# Problems with comparative analyses of avian brain size

**DOI:** 10.1101/2021.11.25.469898

**Authors:** Rebecca Hooper, Becky Brett, Alex Thornton

## Abstract

There are multiple hypotheses for the evolution of cognition. The most prominent hypotheses are the Social Intelligence Hypothesis (SIH) and the Ecological Intelligence Hypothesis (EIH), which are often pitted against one another. These hypotheses tend to be tested using broad-scale comparative studies of brain size, where brain size is used as a proxy of cognitive ability, and various social and/or ecological variables are included as predictors. Here, we test how methodologically robust such analyses are. First, we investigate variation in brain and body size measurements across >1000 species of bird. We demonstrate that there is substantial variation in brain and body size estimates across datasets, indicating that conclusions drawn from comparative brain size models are likely to differ depending on the source of the data. Following this, we subset our data to the Corvides infraorder and interrogate how modelling decisions impact results. We show that model results change substantially depending on variable inclusion, source and classification. Indeed, we could have drawn multiple contradictory conclusions about the principal drivers of brain size evolution. These results reflect recent concerns that current methods in comparative brain size studies are not robust. We add our voices to a growing community of researchers suggesting that we move on from using such methods to investigate cognitive evolution. We suggest that a more fruitful way forward is to instead use direct measures of cognitive performance to interrogate why variation in cognition arises within species and between closely related taxa.

## Introduction

The principal drivers of cognitive evolution have been debated for decades (Ashton et al., 2020; Barton, 1996; Dunbar, 1992; Dunbar & Shultz, 2017; Holekamp, 2007; Humphrey, 1976; Jolly, 1966; Rosati, 2017). Researchers often fall into two broad camps, focusing primarily on either social or ecological factors. Briefly, the Social Intelligence Hypothesis (SIH) posits that cognitive evolution is principally driven by the informational challenges of navigating a dynamic social environment, such as the need to track, anticipate and respond to the behaviour of social partners, and monitor the relationships of others (Byrne & Whiten, 1988; Dunbar, 1998; Humphrey, 1976; Jolly, 1966). In contrast, the Ecological Intelligence Hypothesis (EIH) and its variants emphasise informational challenges posed by ecological variables, such as variable food sources and climatic conditions (Allman et al., 1993; Barton, 1996; Clutton-Brock & Harvey, 1980; Deaner et al., 2003; Harvey & Krebs, 1990). A large body of research has investigated social and ecological correlates of brain size (Chen et al., 2021; DeCasien et al., 2017; Dunbar, 1992; Iwaniuk, 2004; MacLean et al., 2009; Pérez-Barbería et al., 2007; Sayol et al., 2016; Shultz & Dunbar, 2007; Street et al., 2017; van Woerden et al., 2010; West, 2014), but results are often inconsistent and contradictory (Healy & Rowe, 2007; Iwaniuk, 2004; Logan et al., 2018; Powell et al., 2017; Wartel et al., 2019). For instance, Dunbar (1992), found that primate social group size positively correlated with a measure of brain size, which is commonly used as a proxy for cognitive ability. In contrast, DeCasien et al. (2017) found that diet is an important driver of primate brain size but social group size is not. Wartel et al. (2019), on the other hand, found that either diet or social group size could predict primate brain size, depending on where brain size data and predictor variables were sourced from. While most research interrogating the SIH and EIH has focused on primates, birds have emerged as a major model system in cognitive evolution over the last 20 years (Güntürkün & Bugnyar, 2016; Iwaniuk & Arnold, 2004; Overington et al., 2009; Sayol et al., 2016; Seed et al., 2009). Some species of bird show convergent cognitive performance to primates (Güntürkün & Bugnyar, 2016; Seed et al., 2009), yet birds have divergent neuroanatomy (Güntürkün & Bugnyar, 2016) and differing constraints on brain size, such as those imposed by long-range migration (Vincze, 2016). Here we interrogate the potential pitfalls that arise in the comparative study of cognitive evolution in birds. Moreover, we highlight potential pitfalls of current methodologies that have not yet been investigated in any taxa.

The relationship between brain size and cognitive ability is largely unknown and highly contentious (Chittka & Niven, 2009; Healy & Rowe, 2007; Logan et al., 2018). Nevertheless, studies investigating comparative cognitive evolution very often use some measure of brain size as a proxy of cognitive ability (Wartel et al., 2019). Most comparative studies of brain size use a single measurement of brain size per species, taken either from a single “type specimen” individual or averaged across multiple individuals. The influence of intra-specific variation in brain size on the results of comparative analyses are therefore poorly understood. Moreover, to control for the allometric relationship between brain and body size, most studies of brain size control for body size (‘relative brain size’) (Logan et al., 2018). However, the impact that intraspecific variation in body size estimates may have on analyses of relative brain size has not been investigated. Here, we collate data from multiple datasets of brain and body size to interrogate how estimates vary across datasets. Given that most comparative studies of brain size use only one estimate of brain and body size, variation in estimates has the potential to substantially change results.

The approach that researchers take toward model specification may also considerably influence results. While most studies utilise similar statistical techniques to test hypotheses of brain size evolution, approaches toward model specification differ. Indeed, some researchers opt to include covariates associated only with the hypothesis of interest (broadly, the SIH or EIH), and either omit (e.g., Emery et al., 2007) or include less detailed (e.g. Chen et al., 2021; Sayol et al., 2016) variables associated with the competing hypothesis. However, the combination of variables is known to have a substantial influence on the results of primate brain size models (Powell et al., 2017; Wartel et al., 2019). In addition, where covariates are sourced from can have a substantial impact on results. For instance, Wartel et al. (2019) showed that changing the source from which covariates (e.g. diet) were collected substantially changed the results of a previous study (DeCasien et al., 2017). Moreover, decisions on how to define variables can be a somewhat subjective decision made by authors, and this may also have a significant influence on results. For example, if some populations of a species are migratory, but most are resident, should the species be classified as migrant, resident, or a different category altogether, in which case small sample sizes per category may become a concern? How such classification decisions influence model results is, as yet, unquantified.

Here, we use multiple datasets of brain and body size to quantify variation in estimates for more than 1000 species of bird. Following this, we interrogate whether conclusions drawn from models testing alternative hypotheses for brain size evolution differed depending on the combination, source and classification of variables included. To do this, we collated detailed social and ecological variables for species in the Corvides infraorder. The Corvides are well-suited to this investigation because they have a well-resolved phylogeny (Jønsson et al., 2016), large variation in brain size (Iwaniuk & Arnold, 2004), and the majority of species have known social and ecological variables. In addition, focusing on a group of closely related taxa helps to mitigate the potentially confounding effect of differing brain and body size allometries between distantly related taxa (Ksepka et al., 2020; Logan et al., 2018). Together, these investigations allow us to (i) identify novel pitfalls in the study of comparative cognition, and (ii) highlight parallel pitfalls to those previously identified in the field of primate comparative cognition (Powell et al., 2017; Wartel et al., 2019).

## Methods

### Quantifying intraspecific variation in brain and body size

Whole brain volumes across bird species were collated from six published datasets (Corfield et al., 2013; García-Peña et al., 2013; Iwaniuk et al., 2005; Iwaniuk & Arnold, 2004; Iwaniuk & Nelson, 2003; Sayol et al., 2016), all of which recorded brain volume using either the endocranial volume technique (see Iwaniuk & Nelson, 2002 for details), brain mass converted to volume (Iwaniuk & Nelson, 2002), or both. These datasets are non-independent, with some measurements shared between them. Each dataset reported one datapoint per species, with the exception of García-Peña et al. (2013) who reported two datapoints per species (one female, one male) and their associated standard errors (where more than one type specimen per sex was used).

Body sizes were collated from eleven published datasets from ten studies (Corfield et al., 2013; Fristoe et al., 2017; Garamszegi et al., 2002; Iwaniuk et al., 2004, 2005; Iwaniuk & Arnold, 2004; Lendvai et al., 2013; Minias & Podlaszczuk, 2017; Sayol et al., 2016; Sol et al., 2010). One study collated two independently collected sources of body mass data (Minias & Podlaszczuk, 2017) and these were thus treated as two different datasets. All body sizes were measured in grams. Again, datasets are not independent, with some using overlapping sources. Each dataset had one datapoint per species; one dataset recorded standard errors associated with the estimate (Garamszegi et al., 2002).

### Influence of variable inclusion, classification and source

We tested whether including detailed ecological *and* social covariates, relative to including ecological *or* social covariates, qualitatively changed conclusions of models. To do this, we extracted/collated detailed ecological and social variables that have previously been shown to have a significant relationship with brain size (see *Methods: Variables*). In addition to constructing models with differing sets of predictor variables, we examined how sensitive model results were to choices regarding the classification of variables (see *Methods: Variables: Re-classification*). We also tested whether collecting variables from differing sources changed model results (see *Methods: Variables: Environmental variation*).

### Variables

We extracted/collated the following detailed ecological and social variables that have previously been shown to have a significant relationship with brain size.

#### 1. Ecological variables

We included species movement, environmental variability and diet. Species that migrate are thought to have smaller brains than resident species (Pravosudov et al., 2007; Shultz & Dunbar, 2010; Sol et al., 2010; Vincze, 2016). This is hypothesised to be because the energetic cost of the brain constrains selection on increased brain size in migrating species, who have large energetic demands during migration (Pravosudov et al., 2007; Sol et al., 2010; Vincze, 2016). Meanwhile, species that live in fluctuating environments (Fristoe et al., 2017; Sayol et al., 2016; Schuck-paim et al., 2008) and species with broader diets (Sayol et al., 2016) are thought to have bigger brains than those in more stable environments or with specialist diets. This is potentially because species that encounter more ecological uncertainty must process more information in order to respond appropriately (Dall et al., 2005; Sayol et al., 2016; Schmidt et al., 2010), and therefore require more ‘processing power’ (i.e., bigger brains, assuming bigger brains to be a proxy of better cognitive ability).

##### 1.1 Movement

We coded species movement using four categories: resident, partial migrant, migrant or nomadic. Previous studies including migration as a covariate tend to include migration as a binary variable (resident or migratory: Fristoe et al., 2017; Shultz & Dunbar, 2010). However, some species are only migratory in certain regions. Such species were therefore coded as partial migrants (but see *Methods: Variables: Re-classification)*. Meanwhile, other species are neither migrants nor residents and can instead be considered nomadic.

##### 1.2 Environmental variability

We collected environmental variability from two sources. The first measure of environmental variability was ‘temperature variation’, as reported in Fristoe et al., 2017, where higher values indicate more variability. The second was a measure of environmental variability calculated by Sayol et al. (2016). Briefly, Sayol et al. included multiple environmental variables in a phylogenetic principal component analysis. The resultant phylogenetic principal component 1 (PPC1) captured seasonal variation, duration of snow cover and among-year variation, with higher values indicating higher variation, longer snow-cover and larger among-year variation. PPC1 can therefore be interpreted as an axis describing general environmental variation. Meanwhile, phylogenetic principal component 2 (PPC2) captured environmental variation at lower latitudes (e.g., drought events). Temperature variation and PPCs were never used in the same models; instead, they were interpreted as two independent sources of ‘environmental variation’, which we used to quantify whether differing variable source may influence results.

##### 1.3 Diet breadth

We used diet breadth as reported in Sayol et al. (2016) who used Rao’s quadratic entropy (de Cáceres et al., 2011) with diet frequency for seven diet types. Higher values indicate a broader diet.

#### 2. Social variables

We used two social variables in our models, both of which have been suggested to be involved in brain size evolution: social foraging and cooperative breeding. While long-term monogamy has been shown to positively correlate with brain size in some studies (Emery et al., 2007; Shultz & Dunbar, 2010), almost all species in our sample form long-term monogamous pair bonds (see Supplementary Data) so there was not enough variation for this variable to be included.

##### 2.1 Social foraging

Foraging group structure has previously been shown to correlate with relative brain size (Emery et al., 2007; Shultz & Dunbar, 2010). Specifically, species that forage in pairs or bonded groups have been shown to have larger brains than those that forage in large aggregations (Shultz & Dunbar, 2010). Similarly, species that live in small groups have been shown to have bigger brains than those that live in large aggregations (Emery et al., 2007). This is argued to be because the *quality* rather than *quantity* of social bonds is a key driver of cognitive evolution in birds (Emery et al., 2007; Shultz & Dunbar, 2010). However, in other studies foraging group structure appears to be unimportant (Sayol et al., 2016). A common problem with the inclusion of social variables in comparative studies is that they may not capture the underlying informational demands which, according to the SIH, drive cognitive evolution (Boucherie et al., 2019; Dunbar, 1998; Lukas & Clutton-Brock, 2018). We therefore expanded on previous categorisations of foraging group structure by trying to capture variables thought to be associated with information-processing. Specifically, species were coded as foraging solitarily, in pairs, in small groups (<30 individuals), in aggregations (>30 individuals), or as nested versions of these variables (e.g., forages in pairs nested within larger groups). If a species is known to forage in different social contexts but not necessarily in a nested fashion, we categorised these species using the largest group size commonly recorded (e.g., if the species forages in pairs and in small groups, but not necessarily in a nested manner, we recorded this as small group foraging). Following predictions of the SIH, we expected that solitary foragers would have the smallest brains, given the relatively limited demands for processing social information. Moreover, we expected that species foraging in nested groups would generally have larger brains, given the informational demands of managing relationships within a multi-layered context (e.g., managing the pair-bond relationship within a wider social group).

##### 2.2 Cooperative breeding

The role of cooperative breeding in cognitive evolution is contentious. Some authors argue that cooperative breeding entails substantial cognitive demands because individuals need to cooperate and coordinate with multiple others to raise offspring (Burkart et al., 2009; Burkart & van Schaik, 2009; Hrdy, 2009). Conversely, others suggest that the typically high levels of relatedness and shared interests within cooperatively breeding groups may in fact reduce cognitive demands relative to independent breeding (Lukas & Clutton-Brock, 2018; Thornton et al., 2016; Thornton & McAuliffe, 2015). Relevant empirical evidence remains limited and controversial. For instance, Burkart & van Schaik (2009) suggest that cooperatively breeding monkeys show elevated socio-cognitive performance, but these species also have particularly small brains (Thornton & McAuliffe, 2015), and rank poorly in meta-analyses of cognitive performance across primates (Deaner et al., 2006). Among birds, the only comparative study to date found no relationship between cooperative breeding and brain size (Iwaniuk & Arnold, 2004), but this study did not include variables since shown to be significantly related to brain size, such as diet and environmental variation (Sayol et al., 2016). We therefore included cooperative breeding as a binary variable in our analyses. We note that species such as American crows (*Corvus brachyrhynchos*) and carrion crows (*Corvus corone*) are facultative cooperative breeders, but as there were few species in our sample that could be defined as such, we classified all facultative cooperative breeders as cooperative (but see *Methods: Variables: Re-classification)*.

###### Re-classification

Some classifications are ambiguous and multiple different classifications can be justified. We therefore tested whether re-classifying variables changed model results. We re-classified one ecological and one social variable. “Partial migrants”, where at least one population of a species migrates but other populations are resident, were re-classified as residents. Facultative or suspected cooperative breeders were re-classified as non-cooperative breeders, rather than cooperative breeders.

### Statistical modelling

All statistical analyses were undertaken in R v4.0.2 (R Core Team 2017). We used a phylogenetic generalized least squares (PGLS) modelling framework (Freckleton et al., 2002) in the package *caper* (Orme, 2018), which controls for non-independence of datapoints due to relatedness. We used this method because it is the most commonly used technique in the comparative brain size literature (e.g. Fristoe et al., 2017; Fristoe & Botero, 2019; Sayol et al., 2016; Shultz & Dunbar, 2010; Sol et al., 2010; Vincze, 2016). We constructed a consensus tree by downloading 1000 equally plausible phylogenetic trees for the species in our sample from www.BirdTree.org (Sayol et al., 2016). We used the Hackett rather than Ericson backbone because it is the most recently constructed; however, differences between backbones are small and they tend to produce consistent results (Rubolini et al., 2015). Using TreeAnnotator in BEAST v1.10.4, a maximum clade credibility consensus tree was built from these equally plausible trees. This tree was then used to control for phylogenetic non-independence in the following PGLS models. Lambda was estimated using Maximum Likelihood. Model diagnostics and variance inflation factor (VIF) were checked to ensure assumptions were met and variables were not unacceptably collinear, respectively.

#### Corvides analysis

Here, we tested how variable combination, source (environmental variability: temperature variation or PPC) and classification (partial migrant/resident; cooperative breeder/non-cooperative breeder) changed conclusions. See Table 1 for a summary of model formulations. We used Sayol et al.’s (2016) brain size data only. We chose to use this dataset because only one method was used to measure brain size, and all body mass data came from the same specimens that brain volume was taken from. It is therefore likely to be the most precise data currently available. Using this data, we built three models: an SIH model (brain size in response to body size, cooperative breeding and social foraging), an EIH model (brain size in response to body size, migration, environmental variability and diet breadth), and a ‘combined’ model with all covariates included. For the EIH model, we tested two sources of environmental variation: temperature variation (Fristoe et al., 2017) and PPCs (Sayol et al., 2016). We used temperature variation as the measure of environmental variation in the combined model because the limited number of species with PPC data available resulted in some social foraging categories having extremely limited sample sizes. We tested ‘initial’ and ‘reclassified’ variables for species movement and cooperative breeding across all models. Brain and body size measurements were log-transformed.

**Table 1.**
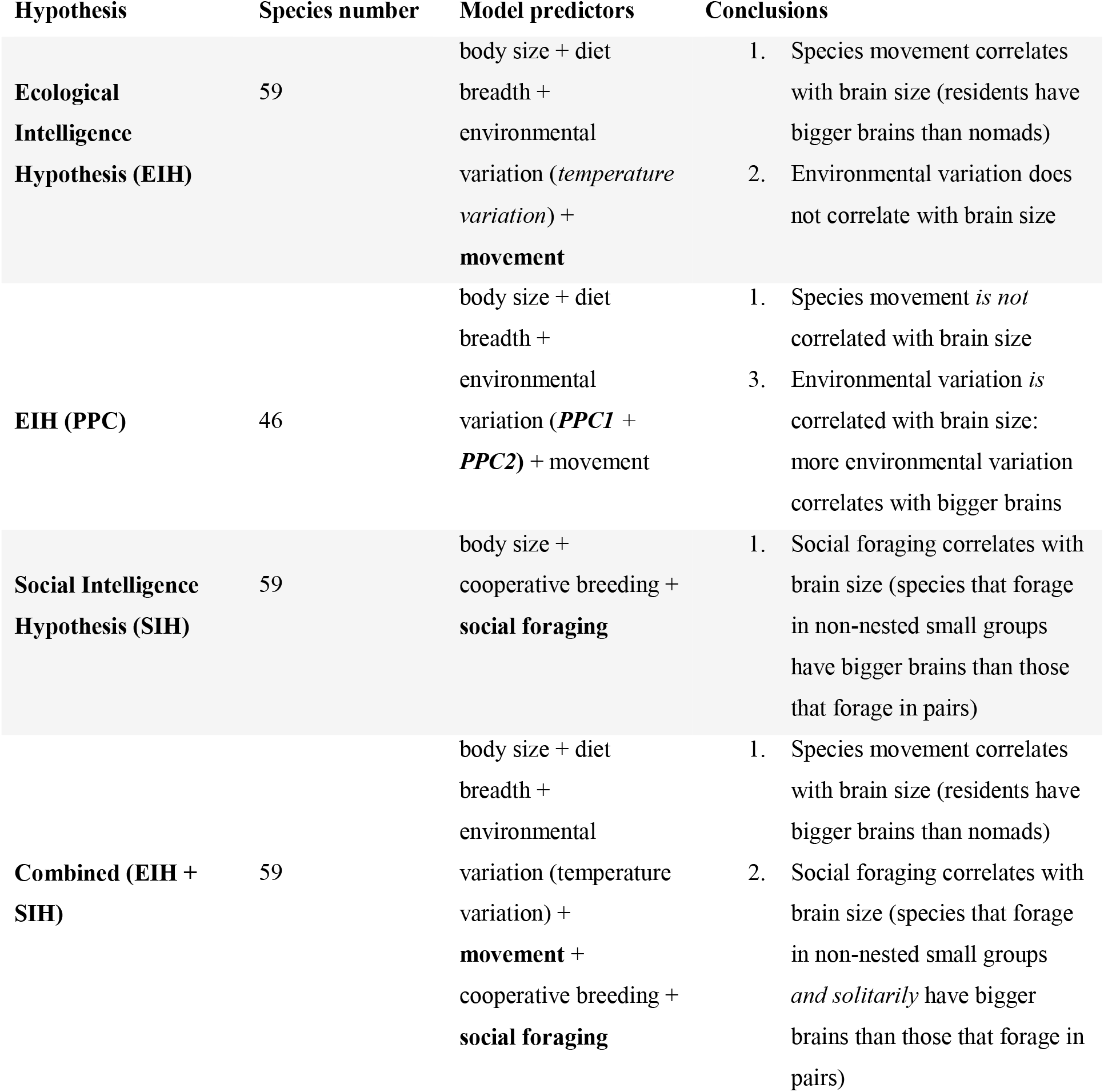
Different models testing a specific hypothesis (SIH/EIH) to test how variable inclusion changes conclusions drawn from results.

## Results

### Quantifying intraspecific variation in brain and body size

Altogether we collated brain size data for 1484 bird species. Of these, 1054 species had brain measurements in more than one dataset. Figure 1a visualises variation in log-transformed brain size estimates across datasets. All but one of the collated datasets had one brain size estimate per species; we therefore present García-Peña’s sex-separated data as sex-averaged in Figure 1a. Only two of the five studies that did not measure sexes separately explicitly stated that brain size datapoints were sex-averaged (Iwaniuk & Arnold, 2004; Sayol et al., 2016). While brain size estimates tended to be the mean value of multiple specimens (on average, ∼six specimens per species), ten or more brain size estimates were based on a single type speciman in at least two studies (Corfield et al., 2013; Iwaniuk et al., 2005).

**Figure 1a.**
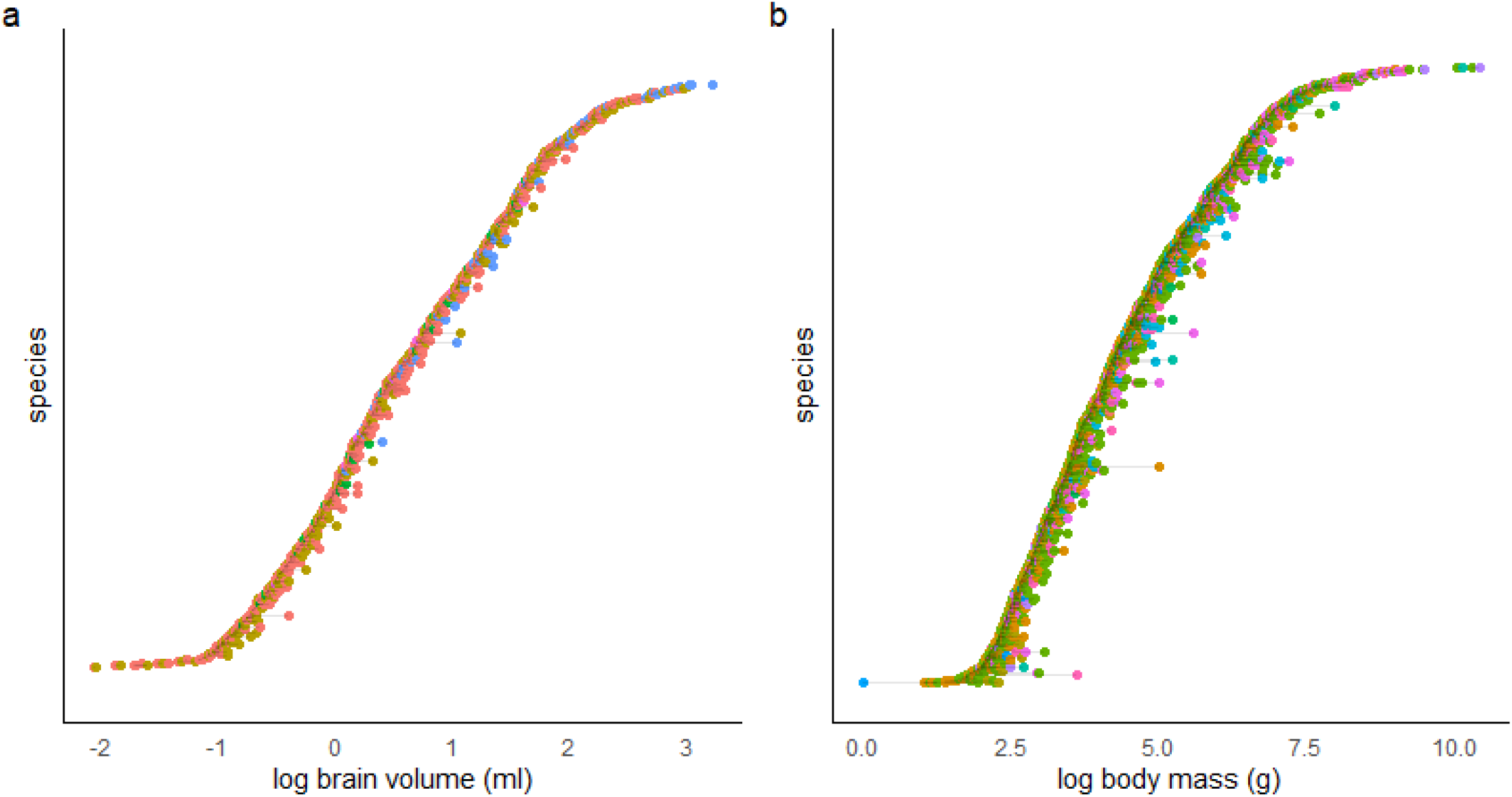
Log brain volume (ml) per species. Only species with brain size estimates in two or more datasets are included 1054). **1b** Log body mass (g) per species. Again, only species with body mass estimates in two or more datasets are included (n = 1578). In both **1a** and **1b**, species are ordered by minimum brain size. Grey lines connect estimates for the same species, which always correspond to estimates from different datasets. Point colour is associated with the source dataset (n = 6 for **1a**, n = 11 for **1b**).

Across datasets, some species varied considerably in brain size estimate (Figure 1a). For instance, the minimum and maximum brain size estimates of *Pyrrhura frontalis* (with the largest overall difference in estimates) overlapped with the brain size estimates of 161 other species in our full sample of 1484 species (10.85%). Similarly, *Acanthorhynchus tenuirostris’* minimum and maximum estimates (with the second largest absolute difference) overlapped with 150 species (10.11% of our sample) and *Tinamus major*’s estimates (the third largest absolute difference) overlapped with 143 species (9.64% of our sample). Variation was not driven by any one dataset in particular (Figure 1a).

Our collated body size dataset contained 2398 species from eleven datasets. 1578 of these species had body mass measurements in more than one dataset. There was a general paucity of details for body mass data. Only one of 11 studies explicitly reported the number of samples from which body mass estimate was calculated (Garamszegi et al., 2002). Only two of 11 explicitly stated that body mass estimates were sex-averaged (Lendvai et al., 2013; Minias & Podlaszczuk, 2017). Ten of the eleven datasets recorded both brain size and body mass data; however, only three of these *sometimes* collected body mass from the same specimens as brain size (Corfield et al., 2013; Iwaniuk & Arnold, 2004; Sol et al., 2010) and only two *always* collected body mass from the same specimens as brain size (Garamszegi et al., 2002; Sayol et al., 2016). Three studies did not mention the primary sources of the body mass data used (Iwaniuk et al., 2004, 2005; Lendvai et al., 2013).

Figure 1b visualises variation in log-transformed body mass estimates across datasets. Again, some species varied considerably in body mass estimates. For instance, the minimum and maximum body mass estimates of *Malurus pulcherrimus* (with the largest overall difference in estimates) overlapped with the body mass estimates of 146 other species in our full sample of 2398 species (6.09%). Similarly, *Alauda arvensis’* minimum and maximum estimates (with the second largest absolute difference) overlapped with 786 species (32.78% of our sample) and *Charadrius tricollaris’* estimates (the third largest absolute difference) overlapped with 781 species (32.57% of our sample). In parallel to the brain size data, variation was not primarily driven by any one dataset (Figure 1b).

### Corvides analysis (Table 1; Table 2; Figure 2)

The full dataset used for this analysis included 59 species, where all species had known social and ecological variables (excluding PPCs). The PPC subset contained 46 species. Conclusions on the principal drivers of brain size evolution in Corvides differed depending on modelling approach (see Table 1 for a general summary; Table 2 and Figure 2 for model results).

**Table 2.** Phylogenetic Generalised Least Squares model results, comparing different model formulations. All significant pairwise contrasts for categorical variables are presented.

| Model type | Predictors | $\lambda$ | Estimate | Standard error | T-value | P-value |
| --- | --- | --- | --- | --- | --- | --- |
| **EIH** | **Body size** | **0.625** | **0.663** | **0.03** | **22.434** | **<0.001** |
|  | Diet breadth |  | 0.184 | 0.377 | 0.487 | 0.629 |
|  | Temperature variation |  | -0.019 | 0.039 | -0.479 | 0.634 |
|  | Movement (partial) |  | 0.133 | 0.086 | 1.542 | 0.129 |
|  | **Movement (resident)** |  | **0.186** | **0.083** | **2.227** | **0.030** |
|  | Movement (migrant) |  | 0.185 | 0.115 | 1.611 | 0.113 |
| **EIH (PPC)** | **Body size** | **0.948** | **0.605** | **0.034** | **17.952** | **<0.001** |
|  | Diet breadth |  | 0.392 | 0.398 | 0.984 | 0.331 |
|  | **PPC1** |  | **0.013** | **0.006** | **2.207** | **0.033** |
|  | **PPC2** |  | **0.031** | **0.015** | **2.072** | **0.045** |
|  | Movement (partial) |  | 0.054 | 0.061 | 0.887 | 0.381 |
|  | Movement (resident) |  | -0.006 | 0.085 | -0.075 | 0.940 |
| **SIH** | **Body size** | **0.698** | **0.668** | **0.029** | **22.685** | **<0.001** |
|  | Cooperative breeding (binary) |  | -0.072 | 0.053 | -1.358 | 0.180 |
|  | Social foraging (nested pairs) |  | 0.079 | 0.061 | 1.292 | 0.202 |
|  | Social foraging (solitary) |  | 0.111 | 0.075 | 1.473 | 0.147 |
|  | **Social foraging (non-nested small groups)** |  | **0.153** | **0.068** | **2.238** | **0.030** |
|  | Social foraging (nested small groups) |  | 0.062 | 0.111 | 0.560 | 0.578 |
| **Combined (EIH + SIH)** | **Body size** | **0.662** | **0.658** | **0.031** | **21.565** | **<0.001** |
|  | Diet breadth |  | 0.161 | 0.389 | 0.414 | 0.680 |
|  | Temperature variation |  | 0.000 | 0.038 | -0.001 | 0.999 |
|  | Movement (partial) |  | 0.070 | 0.863 | 0.812 | 0.421 |
|  | **Movement (resident)** |  | **0.173** | **0.082** | **2.0967** | **0.041** |
|  | Movement (migrant) |  | -0.018 | 0.135 | -0.132 | 0.896 |
|  | Cooperative breeding (binary) |  | -0.079 | 0.055 | -1.430 | 0.159 |
|  | Social foraging (nested pairs) |  | 0.120 | 0.061 | 1.966 | 0.055 |
|  | **Social foraging (solitary)** |  | **0.226** | **0.099** | **2.291** | **0.027** |
|  | **Social foraging (non-nested small groups)** |  | **0.165** | **0.070** | **2.363** | **0.022** |
|  | Social foraging (nested small groups) |  | 0.112 | 0.114 | 0.981 | 0.332 |

**Figure 2.**
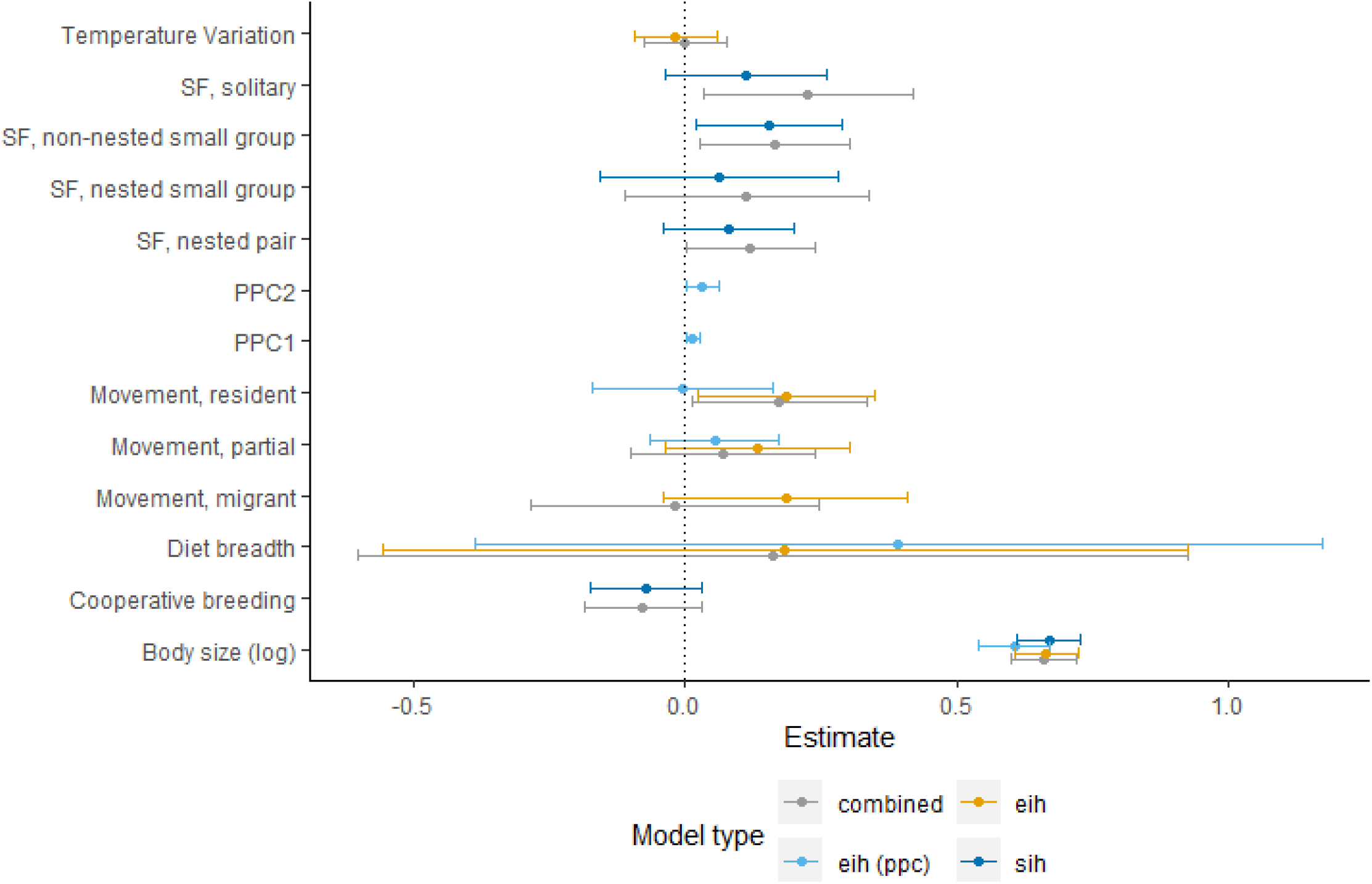
Estimates and confidence intervals for covariates (y axis) generated by models with differing variable combinations (see Table 1). SF = Social Foraging.

In the EIH model with temperature variation included, species movement was significantly associated with brain size. Specifically, resident species were found to have bigger brains than nomadic species. In the EIH model with PPCs (one measure of environmental variation) rather than temperature variation (a different measure of environmental variation) included, both PPC1 and PPC2 were significantly associated with brain size. However, species movement was not. Thus, depending on where we sourced our measure of environmental variation, we could have concluded that environmental variation drives brain size and species movement does not or, indeed, the exact opposite.

Social variables also changed in significance depending on model specification. In the SIH model, species that forage in non-nested small groups were shown to have significantly larger brains than species that forage in pairs. In the combined model, with both social and ecological variables included, solitary foragers and species that forage in non-nested small groups were shown to have significantly larger brains than species that forage in pairs.

Changing facultative or suspected cooperative breeders from their initial categorisation of cooperative breeders to non-cooperative breeders did not qualitatively change SIH model results. However, changing partial migrants to residents did change EIH model results. For the EIH model with PPCs, PPC1 and PPC2 changed from significantly influencing brain size (PPC1: β = 0.01, SE = 0.01; 95% CI[0.001,0.03]; p = 0.03; PPC2: β = 0.03; SE = 0.02; 95% CI[0.002,0.06]; p = 0.04) to having no significant effect (PPC1: β = 0.01, SE = 0.01; 95% CI[0.0001,0.02]; p = 0.055; PPC2: β = 0.03, SE = 0.01; 95% CI[-0.002,0.06]; p = 0.07).

In the combined model, where both changed variables were included, solitary foraging (relative to foraging in pairs) changed from having a significant (β = 0.23, SE = 0.10; 95% CI[0.09,0.37]; p = 0.03) to no significant effect on brain size (β = 0.16, SE = 0.10; 95% CI[-0.03,0.35]; p = 0.11).

Note, however, that although results were unstable in regards to significance, and thus in qualitative conclusions, effect sizes and confidence intervals were not drastically different between models.

## Discussion

In agreement with a growing body of literature (Healy & Rowe, 2007; Logan et al., 2018; Powell et al., 2017; Wartel et al., 2019), our analyses raise concerns that comparative brain size studies are not methodologically robust. We show that there is considerable variation in bird brain and body size estimates across datasets (Figure 1), most likely due to intraspecific variation. The most common methods of comparative brain size analysis do not take this variation into account, although it has the potential to substantially influence results. The combination, source and classification of social and ecological variables also changed results substantially (Figure 2). Indeed, we could have come to several contradictory conclusions depending on our modelling approach. Our results chime with and add to concerns raised in the primate brain size literature (Powell et al., 2017; Wartel et al., 2019) that current methods in the comparative study of brain size evolution give unreliable results.

Comparative brain size studies typically use brain and body sizes averaged from multiple specimens of a single species to obtain one brain and body size estimate per species. Despite all brain sizes in our sample being estimated either from endocranial volume, or brain mass converted to volume (which results in a strong positive correlation with volume measurements (Iwaniuk & Nelson, 2002)), we found substantial variation in brain size estimates across datasets. In several extreme cases, the minimum and maximum brain size measures for one species overlapped with 10% of our sample. Similarly, we found considerable variation between body size estimates of the same species: in extreme cases, the minimum and maximum body size of a species overlapped with more than 30% of the species in our sample. Variation in brain and body size was not driven by any dataset in particular, suggesting that it was not the result of a specific methodological approach, but rather the result of natural intraspecific variation in brain and body size, the use of different specimens by different studies, and the small sample sizes used to obtain estimates. Often, key information about where estimates were derived from was not reported. For instance, the sex of specimens and the variation around the single reported estimate was not reported for most brain size datasets. Body size estimates had even sparser associated information: in most cases, the number of specimens from which the estimate was derived was not given, and neither was sex or variation around the estimate. The fact that estimates tend to be derived from few (or an unknown number of) specimens, often of unrecorded sex, raises concerns as to how well they represent species-average values. Indeed, our study suggests that results of comparative brain size have the potential to be substantially influenced by the brain and body size dataset used, given the variation between estimates.

In addition to investigating variation in brain and body size estimates across datasets, we interrogated how robust models are depending on variable combination, source and classification. Researchers typically have varying approaches to model building, and in agreement with Wartel et al.’s (2019) analyses of primate brain size data, we found that modelling approach influenced results. For instance, depending on whether we chose to include social variables or social and ecological variables, we could have concluded that there is no difference in brain size between species that forage in pairs and species that forage solitarily *or* that species that forage solitarily have significantly larger brains than those that forage in pairs. In addition, we showed that the source of covariates has the potential to substantially change results. Using temperature variation from Fristoe et al. (2017) as a proxy of environmental variation resulted in no support that environmental variation drives the evolution of bigger brains. Meanwhile, using more detailed measures of environmental variation from Sayol et al. (2016) resulted in support. Note, however, that models with temperature variation rather than PPC had a larger sample size, which may influence these results. Nevertheless, these findings parallel those reported in the primate brain size literature (Powell et al., 2017), where using differing variable sources resulted in differing results even when sample sizes were matched. Moreover, we show that variable classification can also drastically influence results. Classifications of variables are sometimes subjective; for instance, species with both cooperatively and non-cooperatively breeding populations could be classified as either. We therefore changed categorical variables that could justifiably be re-classified, and tested how this influenced results. Re-classifying suspected/facultative cooperative breeders as non-cooperative breeders did not change SIH model results; however, re-classifying partial migrants (i.e., where at least one population of a species migrate) as residents substantially changed EIH model results. While two measures of environmental variation (PPC1 and PPC2) were significantly associated with bigger brains before re-classification, there was no significant effect following re-classification. Thus, depending on the classification of a different model covariate, we could have concluded that environmental variation is associated with bigger brains or that there is no association. It must be considered, however, that while p-values crossed the threshold of significance, estimates and confidence intervals for predictor variables did not substantially change. With larger sample sizes, models may be less volatile. Nevertheless, many studies of brain size evolution use sample sizes in the same range as ours (e.g. Schillaci, 2006; Shultz & Dunbar, 2007; Uomini et al., 2020; Vincze, 2016; West, 2014). The concerns raised here are thus pertinent to a wide range of studies.

Taken together, our investigations show that brain and body size data can vary widely between datasets, and that variable combination, source and classification can result in contradictory conclusions depending on somewhat subjective decisions made by researchers. In conjunction with previous work highlighting problems with comparative brain size models (Healy & Rowe, 2007; Logan et al., 2018; Powell et al., 2017; Wartel et al., 2019), this throws previous claims of support for the SIH (e.g., Dunbar, 1992; Shultz & Dunbar, 2010) or the EIH (e.g., DeCasien et al., 2017) into question. In addition to issues with methodological approach, we argue that framing the SIH and EIH as dichotomous and competing hypotheses is not logically sound. We give two reasons for this. First, the hypothesised underlying driver of cognitive evolution for both hypotheses is variation in environmental conditions (including social environment) in which individuals must gather and process information to mitigate uncertainty (Dall et al., 2005; Dunlap & Stephens, 2016). Second, social and ecological variables are not independent, i.e., social species solve ecological problems in a social context, and sociality itself may evolve in response to ecological variables (Ashton, Thornton, et al., 2018; Jetz & Rubenstein, 2011). We therefore suggest not only that our methodological approach to studying comparative brain size evolution needs to change, but also the conceptual framework itself. Rather than splitting often-correlated variables into dichotomous and competing predictors, we could benefit from quantifying the environmental uncertainty animals face in specific contexts and examining how this may drive cognitive evolution.

When considering the accumulating literature on issues associated with comparative studies of brain size evolution (here; Healy & Rowe, 2007; Logan et al., 2018; Powell et al., 2017; Wartel et al., 2019), we add our voices to a growing number in the field suggesting that we move away from such methods to interrogate comparative cognition, and towards a more robust approach that researchers may wish to consider instead. For instance, a fruitful way forward may be to ask how uncertainty in the social and ecological environment influences cognitive performance at the intra-specific level (Ashton, Ridley, et al., 2018; Ashton, Thornton, et al., 2018; Morand-Ferron et al., 2016; Thornton et al., 2014), and between closely related species (Bond et al., 2003; Maclean et al., 2008; MacLean et al., 2013; Sandel et al., 2011; Sheehan & Tibbetts, 2011). Conceptually, we recommend a shift away from treating the SIH and EIH as dichotomous hypotheses; instead, we propose that we should work to understand whether and how uncertainty across a range of contexts drives cognitive evolution.

